# Transcriptional landscape and regulatory roles of small non-coding RNAs in the oxidative stress response of the haloarchaeon *Haloferax volcanii*

**DOI:** 10.1101/218552

**Authors:** Diego R. Gelsinger, Jocelyne DiRuggiero

## Abstract

*Haloarchaea* in their natural environment are exposed to hyper-salinity, intense solar radiation, and desiccation, all of which generate high levels of oxidative stress. Previous work has shown that *Haloarchaea* are an order of magnitude more resistant to oxidative stress than most mesophilic organisms. Despite this resistance, the pathways *Haloarchaea* use to respond to oxidative stress damage are similar to that of non-resistant organisms suggesting that regulatory processes might be key to their robustness. Recently, small non-coding RNAs (sRNAs) were discovered in *Archaea* under a variety of environmental conditions. We report here the transcriptional landscape and functional roles of sRNAs in the regulation of the oxidative stress response of the model haloarchaeon *Haloferax volcanii*. Thousands of sRNAs, both intergenic and antisense, were discovered using strand-specific sRNA-seq, comprising around 30% of the transcriptome during non-challenged and oxidative stress conditions. We identified hundreds of differentially expressed sRNAs in response to hydrogen peroxide induced oxidative stress in *H. volcanii*. Targets of antisense sRNAs decreased in expression when sRNAs were up-regulated indicating that sRNAs are likely playing a negative regulatory role on mRNA targets at the transcript level. Target enrichment of these antisense sRNAs included mRNAs involved in transposons mobility, chemotaxis signaling, peptidase activity, and transcription factors.

**IMPORTANCE:** While a substantial body of experimental work has been done to uncover functions of sRNAs in gene regulation in Bacteria and Eukarya, the functional roles of sRNAs in Archaea are still poorly understood. This study is the first to establish the regulatory effects of sRNAs on mRNAs during the oxidative stress response in the haloarchaeon *Haloferax volcanii*. Our work demonstrates that common principles for the response to a major cellular stress exist across the 3 domains of life while uncovering pathways that might be specific to the Archaea. This work also underscores the relevance of sRNAs in adaptation to extreme environmental conditions.

## INTRODUCTION

Microbial communities that reside inside halite nodules from Salars in the Atacama Desert, Chile, are under extreme environmental pressures due to hyper-salinity, intense solar radiation, and frequent desiccation-hydration cycles, which all generate high levels of oxidative stress.(1, 2) Oxidative stress occurs when the level of reactive oxygen species (ROS) produced in cells overwhelms antioxidant defense mechanisms and damage accumulates.(3) Through metagenomic studies, we found the dominant populations in these salt rocks to be *Haloarchaea* such as *Haloferax* and *Halobacterium*.(4) These halophilic microorganisms are members of the third domain of life, the *Archaea. Haloarchaea* have previously been shown to be highly resistant to ROS damage, withstanding many times what *E. coli* and other radiation-sensitive organisms can survive.(5-7) The haloarcheon *H. salinarum* has been shown to use a wide-range of strategies to combat damage from oxidative stress including multiple copies of genomes (polyploidy) as substrate for DNA repair, functional redundancy of DNA repair and detoxification enzymes (e.g. catalase), increased cytosolic manganese complexes to scavenge ROS, and differential regulation of genes in response to stress.(5-9) However, pathways for DNA repair and protein turnover in *Haloarchaea* are nearly identical to non-resistant bacteria and eukarya suggesting that the regulation of these processes in response to oxidative stress might be key to their robustness. Previous work with *H. salinarum* oxidative stress gene regulatory networks revealed that a single transcription factor, RosR, regulates the appropriate dynamic response of nearly 300 genes to reactive oxygen species stress.(5) This work demonstrated that the oxidative stress response in *H. salinarum* impacted a wide array of cellular processes, engaging at least 50% of all the genes.(2) These results underline the importance of gene regulation in *Haloarchaea* for responding to and counteracting the damage caused by oxidative stress.

Besides transcription factors, small regulatory RNAs (sRNAs) similarly act as global gene regulators.(10) Small RNAs (sRNAs) are ubiquitously found in *Bacteria* and *Eukarya*, playing large-scale roles in gene regulation, transposable element silencing, defense against disease state, and foreign elements.(11-14) Several types of sRNAs have been identified in the *Eukarya* (miRNAs, siRNAs, and piRNAs) and they are typically 20-25 nucleotides (nt) long. Their major mode of interaction is through base pairing to the 3’-unstranslated region (UTR) of their target mRNAs, inhibiting translation or triggering target degradation with associated protein components (Argonautes).(10) Bacterial sRNAs have been shown to modulate core metabolic functions and stress related responses, such as nutrient deprivation, by binding target mRNAs and causing their degradation or preventing translation.(11, 15) Most of the functionally characterized sRNAs in *Bacteria* bind the 5’-UTR of their target mRNA and are longer than their eukaryal counterparts, with sizes ranging from 50 to 500 nt. These sRNAs can target multiple genes, including key transcription factors and regulators. (11, 15, 16) As a consequence, a single sRNA can modulate the expression of large regulons and thus have a significant effect on metabolic processes. For example, the bacterial sRNA OxyS, which is dramatically induced by oxidative stress, regulates the expression of about 40 genes and interacts directly with eight target mRNAs.(11)

sRNAs have been discovered to be abundant in *Archaea*, more specifically in *Haloarchaea*, in response to a variety of environmental conditions but the functional roles of these RNAs still remain poorly understood nor has a link between sRNAs expression and oxidative stress response been established.(13, 17-24) Only a handful of studies on sRNAs in hyperthermophiles, methanogens, and the haloarchaeon *Haloferax volcanii* have been reported so far.(13, 17-24) In *H. volcanii* a large number of intergenic- and antisense-encoded sRNAs, 145 and 45, respectively, were discovered using microarray in addition to a novel class of sRNAs recently described in eukaryotes, tRNA-derived fragments (tRFs), and a new study found thousands of sRNAs present in this organism.(19, 25) In *Sulfolobus solfataricus*, 125 trans-encoded sRNAs and 185 cis-antisense sRNAs were identified using high-throughput sequencing (HTS), suggesting that 6.1% of all genes in *S. solfataricus* are associated with sRNAs.(26) A comparative genome analysis of *Methanosarcina mazei*, *M. bakeri*, and *M. acetivorans* revealed that 30% of the antisense and 21% of the intergenic sRNAs identified were conserved across the 3 species.(27) Co-immuno-precipitation with the Lsm protein (archaeal Hfq homolog) was used to “capture” sRNAs.(17) Some *Archaea* contain eukaryotic Argonaute homologs but their interaction with sRNAs is still yet to be elucidated.(28) All together, these studies suggest that sRNAs are as widespread and abundant in the *Archaea* as in the *Bacteria* and *Eukarya*.

Target mRNA identification of sRNAs has proven to be difficult within the *Archaea* but a necessary task for uncovering sRNA functionality. RNA-seq in *M. mazei* cultures, grown under nitrogen starvation conditions, showed the differential expression of a number of sRNAs in response to nitrogen availability, and allowed for the identification of the first *in vivo* target for archaeal intergenic sRNAs.(27, 29) The potential target for sRNA_162_ is a bicistronic mRNA encoding for a transcription factor involved in regulating the switch between carbon sources and a protein of unknown function.(29) In *Pyrobaculum*, 3 antisense sRNAs were found opposite a ferric uptake regulator, a triose-phosphate isomerase, and transcription factor B, supporting a potential role for archaeal antisense sRNA in the regulation of iron, transcription, and core metabolism.(30) sRNA deletion mutants can be used to identify potential biological functions and target genes. Deletion strains were successfully generated for *H. volcanii*, and phenotyping of the sRNAs deletion mutants revealed several severe growth defects under high temperatures, low salt concentrations, or specific carbon sources.(22, 31) While these studies revealed that sRNAs likely play essential roles in the physiological response to environmental challenges in the *Archaea*, the functional roles and mechanisms of action of these important post-transcriptional regulators still remain unknown. Furthermore, no work has been done to investigate archaeal sRNAs in response to oxidative stress, a universal and frequent stressor in all domains of life that results in extensive cellular damage. In order to determine the impact of sRNAs during the oxidative stress response, we assessed the *H. volcanii* transcriptional landscape during non-challenged and oxidative stress conditions using comparative strand-specific small RNA-sequencing (sRNA-seq).

## RESULTS

To identify globally small non-coding RNAs differentially expressed in response to oxidative stress in *H. volcanii,* we exposed 5 replicate cultures of *H. volcanii* to 2 mM H_2_O_2_, a dose that resulted in the survival of 80% of the cells **(Fig S1)**. RNA from these H_2_O_2_ treated cultures, and from non-challenged cultures (controls), were sequenced using a strand-specific size-selected sRNA library preparation essential for sRNA discovery.

### Small non-coding RNA discovery in *H. volcanii*

We obtained at total of 137 million sequence reads (41 Gb), across all replicates and conditions. Following quality control and reference-based read mapping, we intersected the mapped reads against the *H. volcanii* reference genome to discover sRNA transcripts that we classified as antisense (overlapping a gene and/or its regulatory elements on the opposite strand) **(Fig 1a)** and intergenic (non-coding region between two genes) **(Fig 1b)**. These novel transcripts were 50 to 1,000 nucleotides (nt) in length and represented 35 to 40% of the total transcriptome (**Table S1**). The sRNAs were validated using two in-silico approaches, as described above, and the majority of sRNAs (>90%) passed our validation parameters. Analyzing the upstream regions of sRNAs enabled the discovery that 30% of sRNAs contained both a BRE and TATA-box with centroids at −38 and −29 nucleotides **(Fig. S2)**. Using less conservative parameters (−3, +3 nucleotides) for BRE and TATA-box centroids resulted in 70% of sRNAs containing transcriptional motifs.

**Figure 1:**
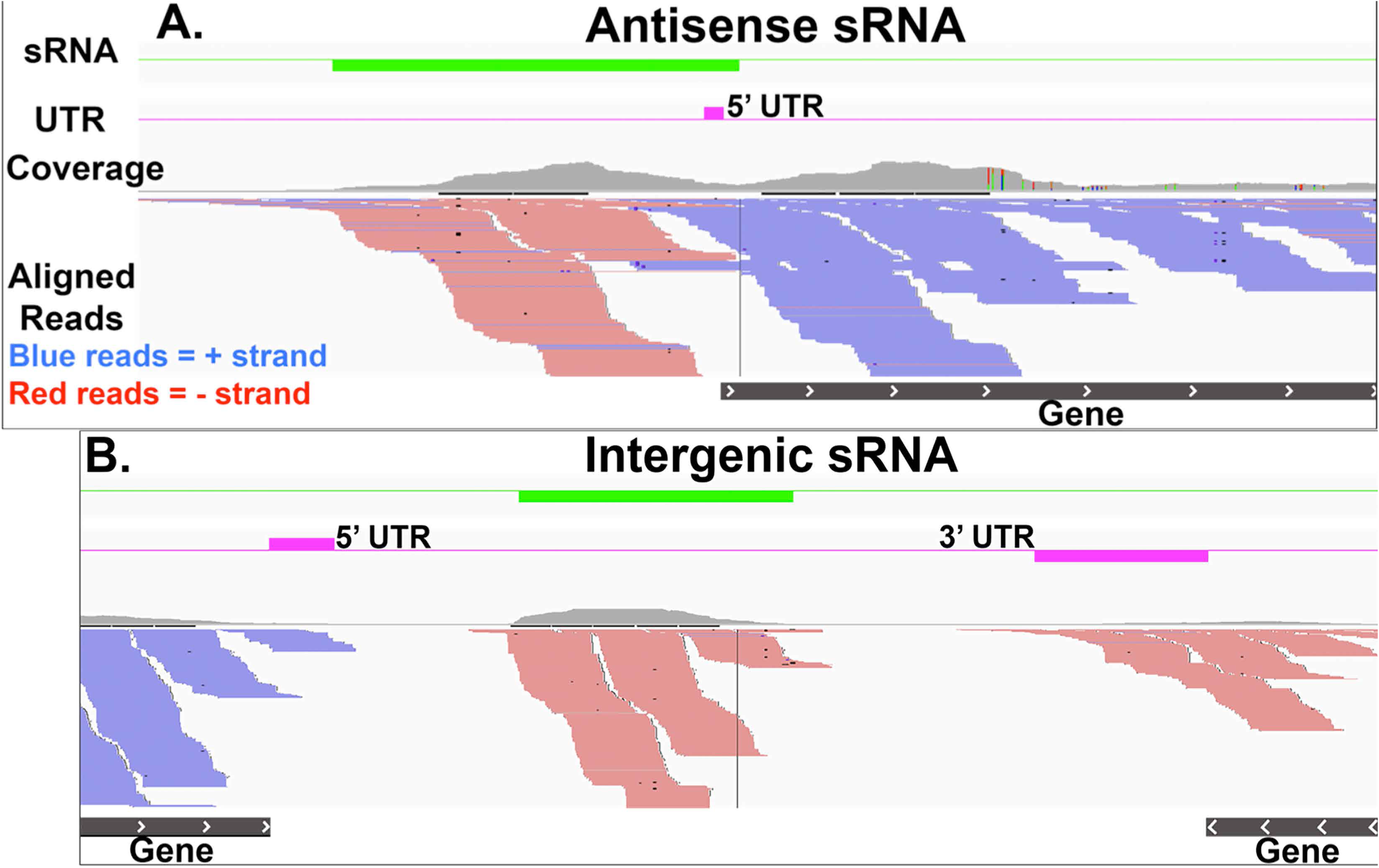
Genome viewer of (A) Antisense sRNAs (*cis*-acting) and (B) Intergenic sRNAs (*trans*-acting). Paired-end reads (100 bases) were mapped to the *H. volcanii* NCBI reference genome. Reference genes are marked as black lines with white arrows indicating their location on the plus strand (>) or minus strand (<). Reads marked in red are transcribed from the minus strand while blue reads are transcribed from the plus strand. Untranslated regions were predicted using Rockhopper2 (pink lines). Green lines mark discovered sRNAs. Coverage plots are in gray.

### Non-coding sRNA characterization in *H. volcanii* during non-challenge conditions

Normalized expression values in RNA-seq analyses are often reported as Reads or Fragments Per Kilobase of transcript per Million mapped reads (RPKM/FPKM). However, RPKM/FPKM have been shown to be inconsistent for comparison between samples (due to transcript length) and another expression value, transcripts per million (TPM), was found preferable for comparison because it is independent of mean expressed transcript length.(32-35) Due to the generally smaller length of sRNA transcripts and variability in size, we chose to use TPM in our analysis to minimize transcript length bias. *H. volcanii* grown under non-challenged conditions (42°C, complex media) expressed a total of 2,577 sRNAs after quality control (transcripts per million (TPM) >0) **(Table S1)**, ranging from 49 to 1,000 nucleotides in size and with an average length of 373 nt. A majority of these sRNAs, 2,493 sRNAs (97%), were antisense to coding-regions **(Fig 2)**. Three of the sRNAs were antisense to CRISPR arrays, suggesting their role in regulating CRISPR systems. The *H. volcanii* H53 auxotroph genome is 4 Mbp and contains 4,130 genes. The genome is comprised of a chromosome stably integrated with plasmid pHV4, 2 plasmids (pHV1, pHV3), and has been cured of plasmid pHV2. A majority of sRNAs (68%) were encoded on the chromosome and integrated plasmid pHV4 (18%). No sRNA encoded on plasmid pHV2 were found, as expected, while sRNAs were encoded on the remaining plasmids pHV1 (2%) and pHV3 (12%).

**Figure 2:**
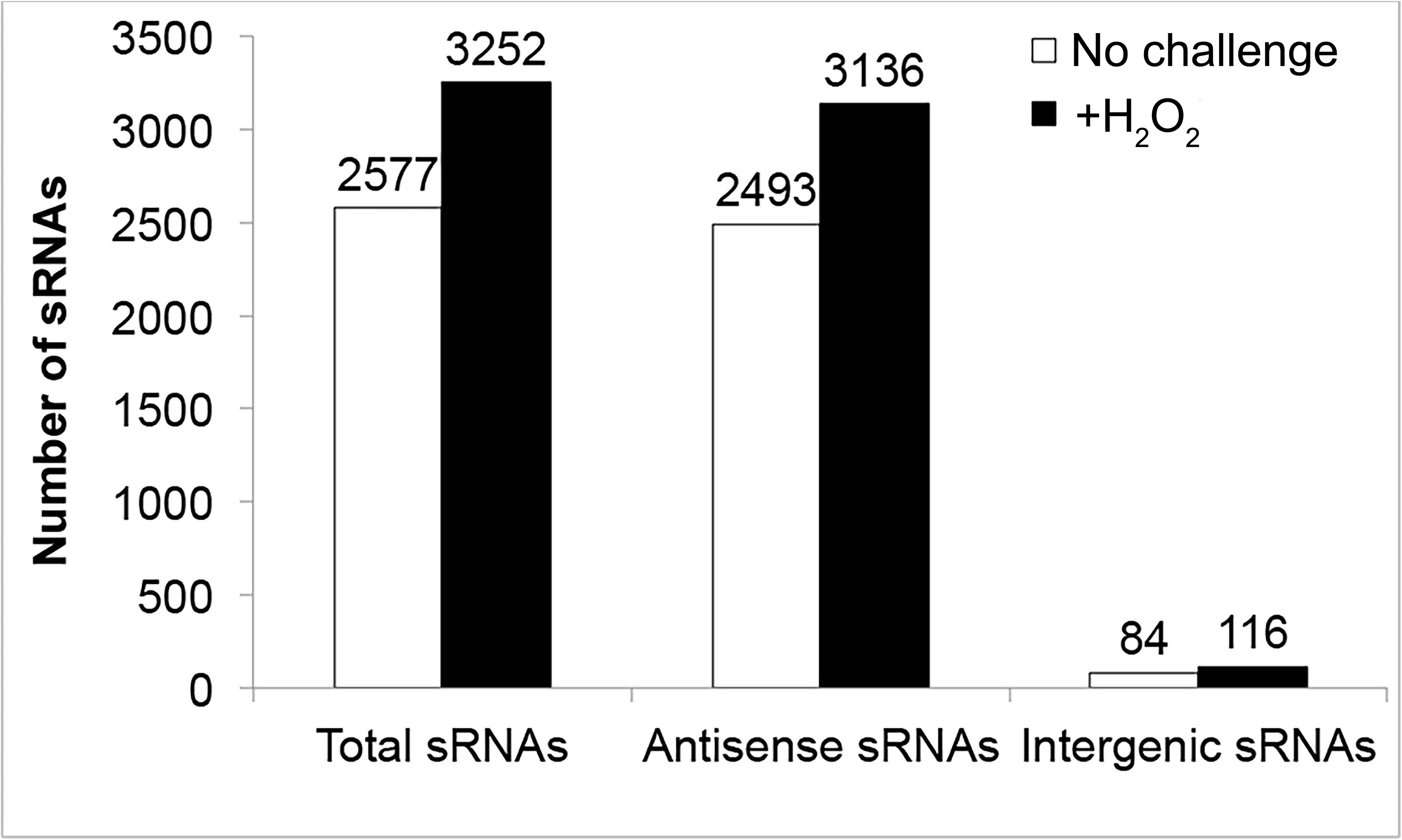
Number of sRNAs (total, antisense, and intergenic) discovered during non-challenged and H_2_O_2_ challenged conditions.

The average expression of the sRNAs was 22.1 TPM. Relative to mRNA expression levels (average: 254.8 TPM), the expression of the sRNAs was on average an order of magnitude lower. A comparison of the distribution of expression levels between sRNAs and mRNAs further confirmed that a majority of sRNAs was more lowly expressed than mRNAs. Of the discovered sRNAs, 75% had expression values less than or equal to 1 TPM **(Figure S3)**, 5% had expression levels similar to that of mRNAs, (TPM >20), and 22 had robust expression levels with TPMs ranging from 100 to 700. Lastly, 2 sRNAs around 150 nucleotides in size exhibited extremely high expression level with TPMs of 14,000 and 460,000, respectively. Transcript length did not correlate with expression levels, indicating that the low expression of sRNAs observed was not an artifact of sequencing (i.e. longer transcripts receiving more read coverage thus skewing coverage based on length) **(Figure S4)**. We found that 4 of the top 5 most highly expressed sRNAs (TPM >150) were located in intergenic regions.

Putative mRNA targets for the most highly expressed antisense sRNAs were identified as the *cis*-mRNA encoded on the opposite strand with a minimum overlap of 25 nucleotides. These targets included an IS4 Family Transposase, ATP—cob(I)alamin adenosyltransferase, glycine dehydrogenase aminomethytransferase, transducer protein Htr36, pyridoxamine 5’-phophase oxidase, XerC/D integrase, deoxyhypusine synthase, and protein translocase TatA. We do not report putative targets for intergenic sRNA because of the inherent difficulty in reliably predicting these targets due to unknown degrees of complementarity (i.e. gaps in hybridization between an intergenic sRNA and a mRNA).

### Non-coding sRNAs in *H. volcanii* during oxidative stress conditions

*H. volcanii* under H_2_O_2_–induced oxidative stress conditions expressed 3,251 sRNAs, a 20% increase in number of sRNAs compared to the non-challenged conditions (**Fig 2, Table S1**). A pattern of sRNA distribution similar to that of the non-challenged condition was observed; more than 90% of sRNAs were antisense and a majority (69%) were encoded on the main chromosome. A smaller average length of 337 nt was observed. Overall TPM expression of sRNAs during oxidative stress was similar to the non-challenged state, with a marked decreased in expression level for the single most highly expressed sRNA (non-challenged: 40926.9 TPM, H_2_O_2_: 16158.6 TPM), which was an intergenic sRNA.

Putative targets for the most highly expressed antisense sRNAs included the 16S rRNA genes (two sRNAs), an IS4 transposase, an MBL fold hydrolase, transducer protein Htr36, pyridoxamine 5’-phosphate oxidase, and a stomatin-prohibitin-like protein. Of the most highly expressed sRNAs, 3 targeted the same mRNAs during both the non-challenged and oxidative stress conditions. These mRNAs encoded for an IS4 family transposase, transducer protein Htr36, and pyridoxamine 5’-phosphate oxidase.

### Regulatory effects and differential expression of sRNAs during oxidative stress

To investigate the regulatory effects sRNAs on their target mRNAs, we compared the expression levels (TPM) of the most highly expressed antisense sRNAs (TPM>50) against the *in silico* determined mRNA targets. We found that the expression of these putative mRNA targets was always lower than that of the sRNA (p ≤ 0.05), for both experimental conditions, with the exception of the sRNAs targeting the 16S rRNA gene (**Fig 3a, Table S1**). This was in contrast to the overall trend in expression levels between sRNAs and mRNAs we reported in **FigS3**, suggesting that the most highly expressed sRNAs may lower the expression of their mRNA targets. When looking at sRNAs with lower expression (10 to 1 TPM) we found that not all but many of these sRNAs had expression equal to or higher than the target mRNA indicating a similar regulatory effect. An example of this negative regulatory effect was seen in the antisense sRNA targeting the IS4 family transposase, which had increased expression during oxidative stress compared to the non-challenged state (H_2_O_2_: 1382.9 TPM, non-challenged: 590 TPM) and, correspondingly, the IS4 family transposase mRNA had decreased expression during oxidative stress (H_2_O_2_: 1.2 TPM, non-challenged 2.7 TPM). The other two sRNAs had increased expression during non-challenged conditions compared to oxidative stress conditions (non-challenged: 369.4; 312.5 TPM, H_2_O_2_: 244.9; 165 TPM) with a similar trend of target mRNA expression decreasing (non-challenged: 3.9; 1.1 TPM, H_2_O_2_: 6.7, 5.4 TPM).

**Figure 3:**
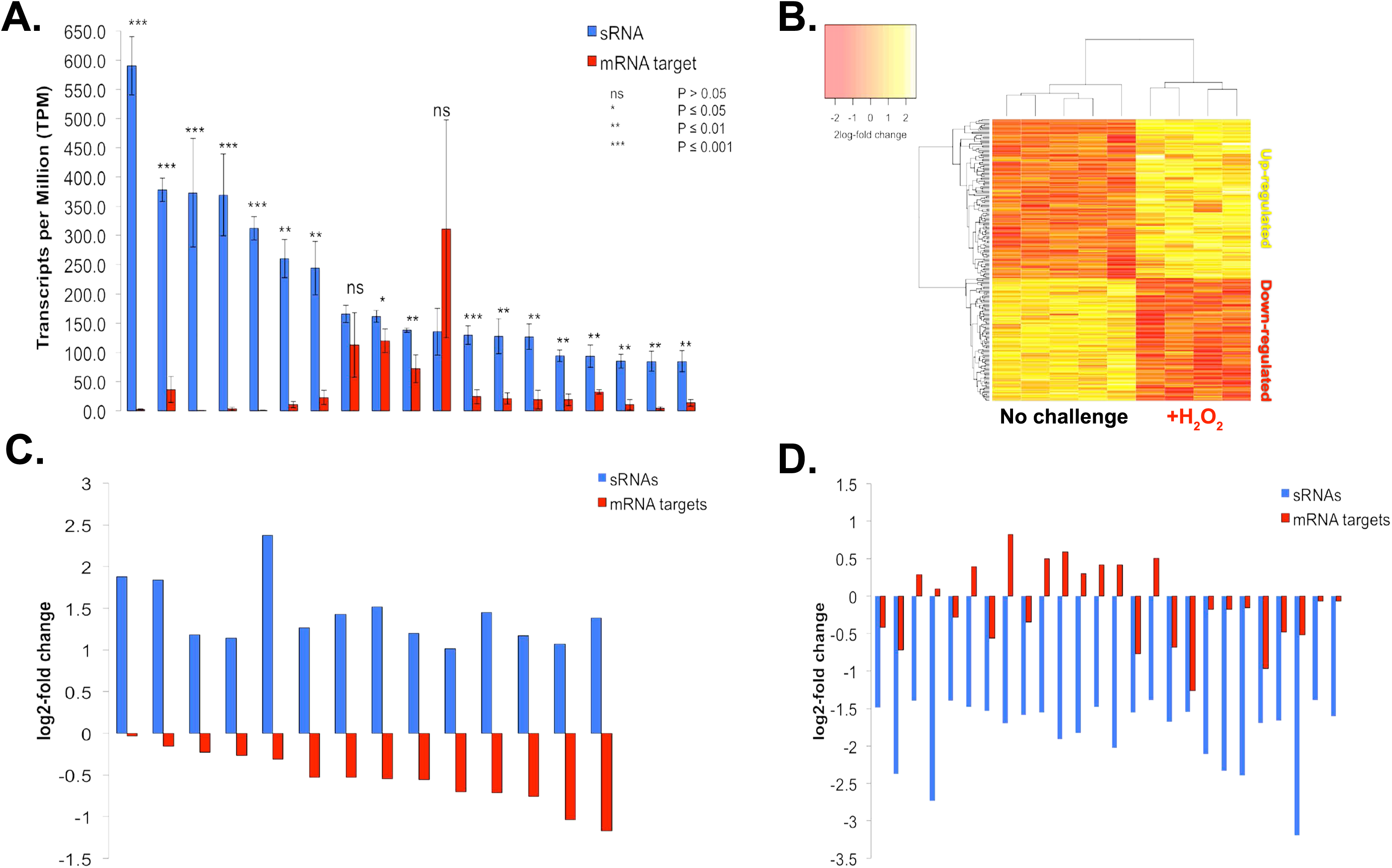
Regulatory effects of differentially expressed sRNAs on their putative mRNA targets during oxidative stress. (A) Expression (TPM) of sRNAs and their respective mRNA targets during oxidative stress. Stars indicate a significant difference in expression between a sRNA and its mRNA target based on a pair-wise student’s t-test; *p-value ≤ 0.05, **p value ≤ 0.01, and *** p-value ≤ 0.001. (B) Heatmap of log transformed fold-change of differentially expressed antisense sRNAs. (C) Differential expression fold changes of up-regulated sRNAs and their mRNA targets. (D) Differential expression fold changes of down-regulated sRNAs and their mRNA targets.

To further investigate this negative regulatory relationship between sRNAs and mRNA targets we probed for differentially expressed sRNAs between the non-challenged and the oxidative stress conditions. Candidate sRNAs were considered significantly up-or down-regulated by oxidative stress using a False Discovery Rate (FDR) of less than 5%. Using this statistical framework, we identified a core set of differentially expressed sRNAs specific to oxidative stress. Both intergenic and antisense sRNAs were differentially expressed. Of the intergenic sRNAs, 55 were significantly differentially expressed, with 27 up-regulated and 28 down-regulated **(Fig S5, Table S2)**. All up-regulated intergenic sRNAs had greater than or equal to 1.3 log_2_-fold change increase in expression during oxidative stress, with the most up-regulated intergenic sRNA having a 4.3 log_2_-fold change increase. A few down-regulated intergenic sRNAs had small fold changes in expression (<1 log_2_-fold change) but most exhibited robust down-regulation (-3.2 log_2_-fold change). A total of 274 antisense sRNAs were either up-regulated and down-regulated during oxidative stress, indicating two populations of antisense sRNAs (**Fig 3b, Table S3**). Seventeen percent (46 sRNAs) of these differentially expressed sRNAs demonstrated a fold-change in expression of 2 or greater; the most up-regulated sRNA had a log_2_-fold change of 5.3 and the most down-regulated sRNA had a log_2_-fold change of −3.6, indicating a role in the cellular response to oxidative stress. Twice the number of antisense sRNAs were up-regulated with a fold-change in expression of 2 or greater (31) compared to down-regulated (15).We then compared differential expression levels between antisense sRNAs and their putative mRNA targets (no putative targets could be reliably identified *in silico* for intergenic sRNAs) and found that, in most instances, up-regulated antisense sRNAs had putative mRNA targets that were down-regulated during oxidative stress (**Fig 3b**). For example, during oxidative stress,14 up-regulated antisense sRNAs targeted transposase mRNAs and each of the cognate transposase mRNAs were found to be down-regulated (**Fig 3c**). Furthermore, only a small subset of down-regulated antisense sRNAs had their mRNA target up-regulated during oxidative stress, while most of the mRNA targets were also down-regulated (**Fig 3b and d**).

Oxidative-stress responsive antisense sRNAs were bioinformatically predicted to overlap both the 5’ and 3’ UTRs of mRNAs indicating a hybrid system between Eukarya (3’ UTR-binding) and Bacteria (5’ UTR-binding) sRNA regulatory systems (**Fig 4**). We found that 7% of antisense sRNAs overlap at the 5’ UTR and 26% overlapping at the 3’ UTR. However, the majority of the antisense sRNAs (67%) were found to overlap the coding sequence (CDS) of mRNAs rather than targeting the UTRs, which has not been previously reported (**Fig 4**). Using Northern blots, we recapitulated the *in vivo* differential expression patterns of selected candidate sRNAs, further confirming transcript size and differential expression levels for oxidative stress, even for the most lowly expressed sRNA candidate (1 TPM) (**Fig 5a and 5b**). We also showed that the strandedness (the strand on which the sRNA was encoded) predicted by our sRNA-seq analysis was confirmed by our *in vivo* data using oligo probe northern blotting of 5’ UTR, 3’ UTR, CDS antisense sRNAs, and intergenic sRNAs (**Fig 5b**).

**Figure 4:**
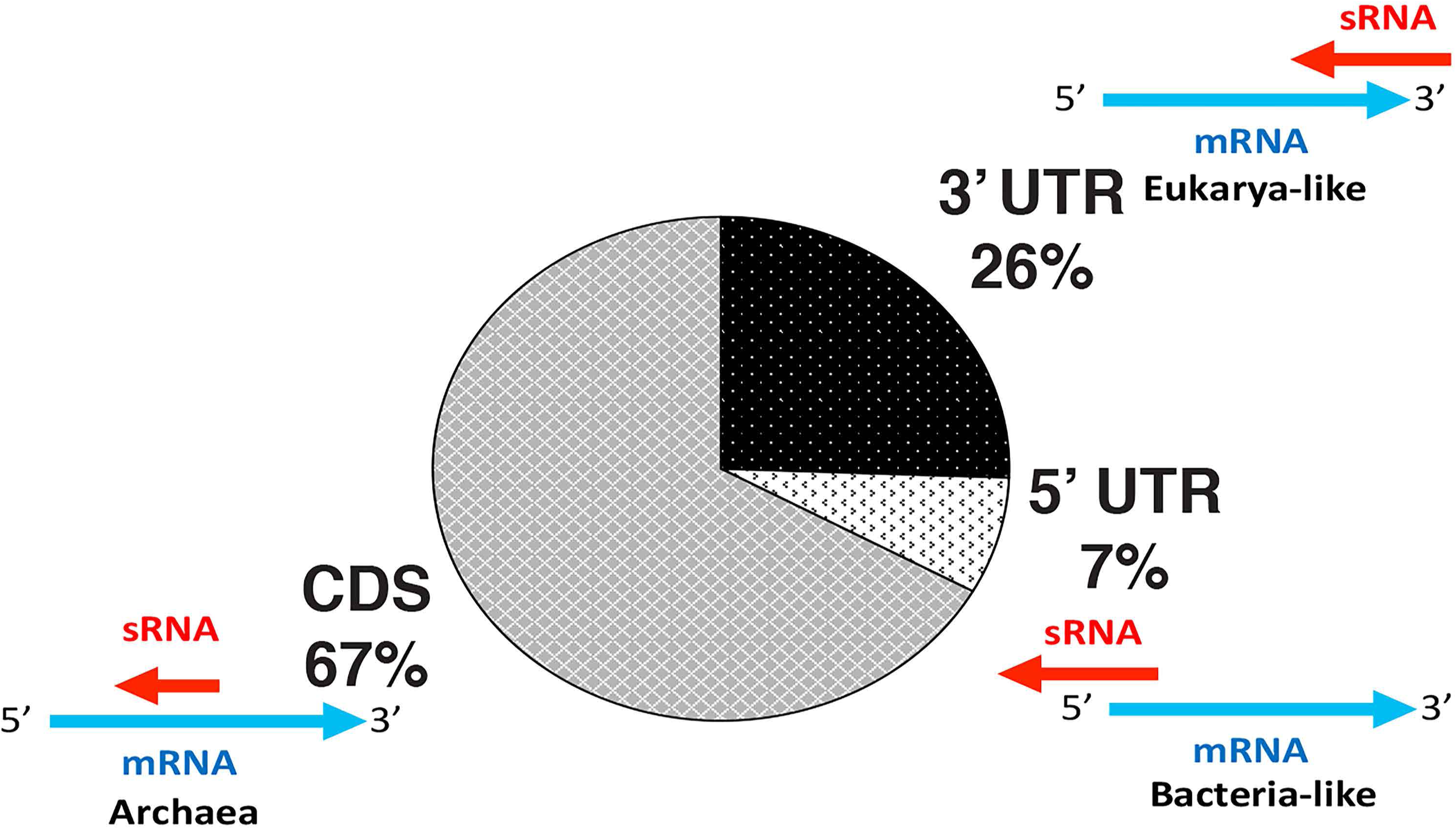
Distribution of binding regions for antisense sRNAs. UTR, untranslated region; CDS, coding sequence.

**Figure 5:**
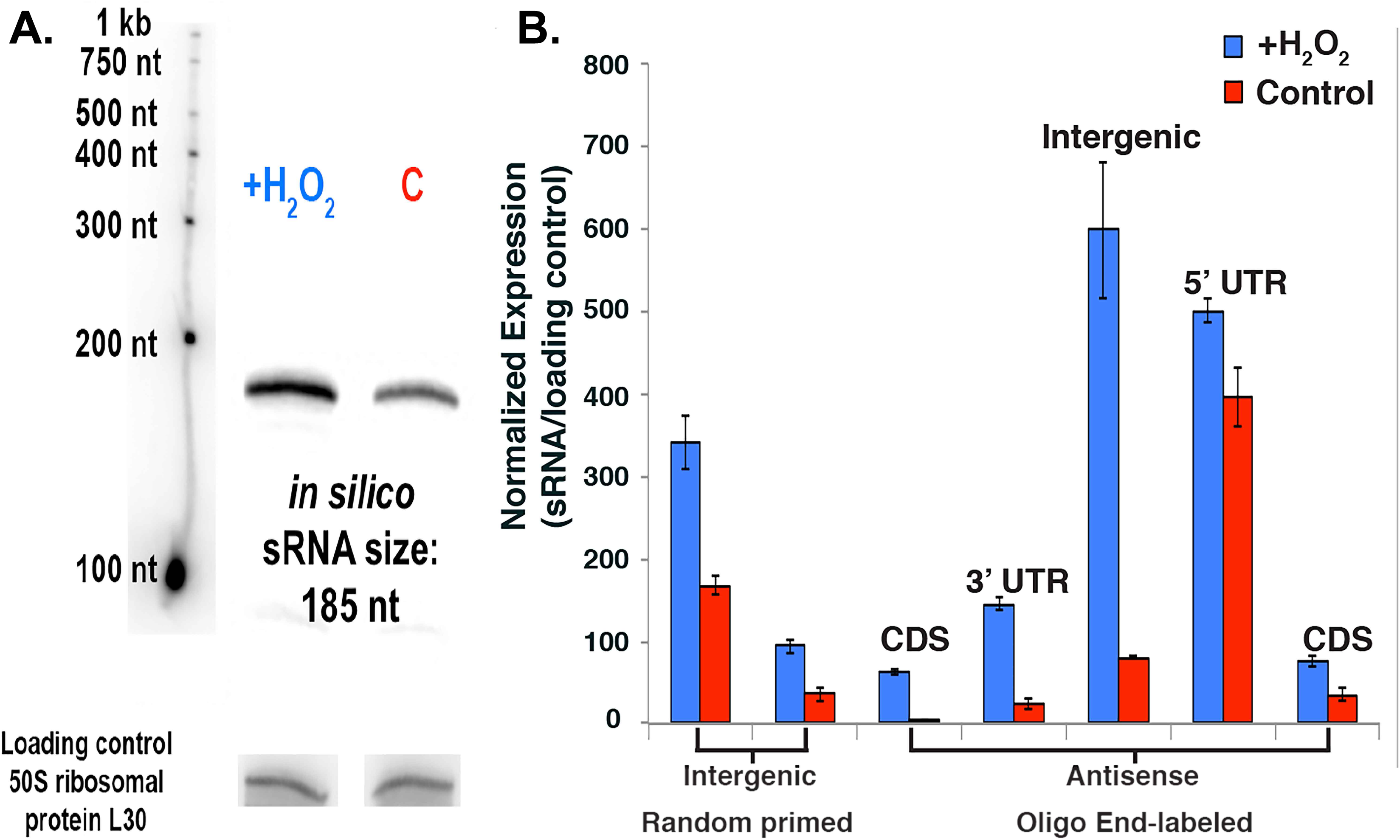
Validation of differentially expressed sRNAs by Northern blots. (A) Representative Northern blot confirming size and differential expression patterns of an intergenic sRNA during oxidative stress. (B) Quantification of Northern blots confirming the expression of lowly expressed sRNAs (random primed labeling) and strand-specificity of sRNAs (oligo labeling). All classes of sRNAs were confirmed: antisense (5’ UTR, 3’ UTR, CDS) and intergenic sRNAs.

### Target Enrichment of sRNAs

We identified *in silico* targets for the differentially expressed oxidative-stress responsive antisense sRNAs. Genes encoded by the putative target mRNAs were categorized by cellular function using the **ar**chaeal **C**luster of **O**rthologous **G**enes (arCOGs) and by pathways using gene ontologies (GO) from the **D**atabase for **A**nnotation, **V**isualization and **I**ntegrated Discovery (DAVID). For sRNAs up-regulated during H_2_O_2_ stress, we found a functional enrichment of target genes encoding transposases, involved in chemotaxis methyl-receptor signaling and in transcriptional regulation (transcription factors) (p <0.05). Genes, from many other pathways that were not enriched, were also the target of antisense sRNAs, including peptidase activity genes and serine and threonine biosynthesis genes (**Fig 6a**). Twenty three of these sRNAs targeted transposase genes. Each transposase gene was down-regulated while their cognate sRNA was up-regulated, and the sRNA was always located at the 5’ UTR of its target. Most transposases belonged to the IS family of transposases except for one DDE transposase. Three transcription factor families (IclR, ArcR, and Asn[C]) were also targeted by antisense sRNAs. A functional enrichment gene ontology analysis found that down-regulated sRNAs target genes were involved in membrane transport (ABC) transporters and biosynthesis of secondary metabolites, as well as targeting hydrolases (**Fig 6b**). A significant proportion of enriched targets for both up-and down-regulated sRNAs were genes encoding hypothetical proteins.

**Figure 6:**
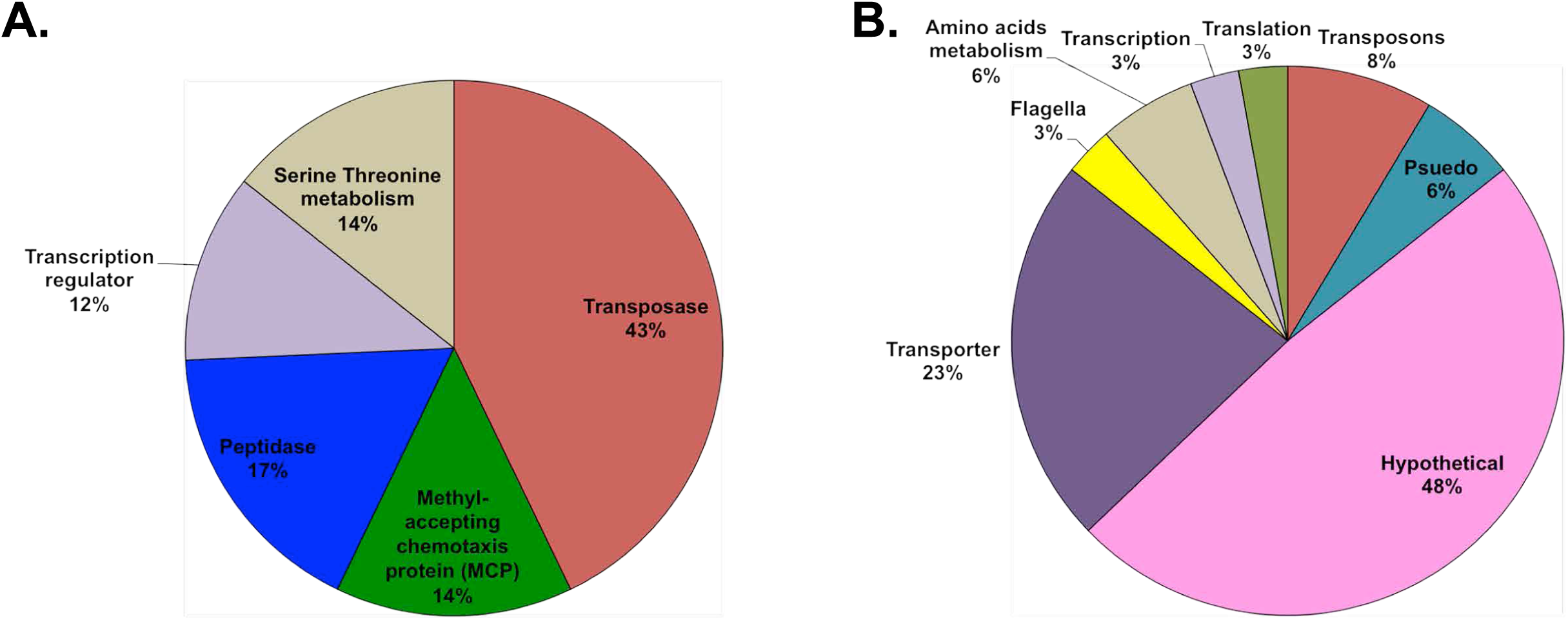
Gene ontology enrichment analysis identifying the functional classification of gene targets of sRNAs during oxidative stress. (A) Enriched target gene functions for up-regulated sRNAs. (B) Enriched target gene functions for down-regulated sRNAs.

### mRNA transcriptional response to oxidative stress in *H. volcanii*

To determine the transcriptional landscape of mRNAs during oxidative stress, especially for mRNAs that were predicted targets of sRNAs, we sequenced rRNA-depleted mRNA-seq libraries in parallel with the previously described sRNA-seq libraries (derived from the same pool of total RNA). During H_2_O_2_-induced oxidative stress, a fourth of all genes (1,176) were significantly differentially expressed with a False Discover Rate less than 5% (**Table S4**). Both catalase and superoxide dismutase, known ROS detoxification enzymes, were up-regulated at the mRNA level thus validating our experimental approach and characterizing *H. volcanii* response to oxidative stress at the transcriptional level (**Fig 7**). A GO enrichment analysis (DAVID) was used to identify what pathways were enriched with differentially expressed genes during oxidative stress. The most enriched (p<0.05) up-regulated genes were involved in transcription, including various transcription factor families, all of the RNA polymerase subunit genes, and transcription initiation factors. Other enriched (p<0.05) up-regulated pathways were involved in iron-sulfur cluster assembly, DNA topological change (topoisomerase), proteasome, cell redox homeostasis, histidine metabolism, and 2-oxocarboxylic acid metabolism. The most up-regulated gene was a reactive intermediate/imine deaminase with a log_2_-fold expression increase of 6.4. The most enriched (p<0.05) down-regulated genes were Tn5-like IS4 transposases. Other enriched (p<0.05) down-regulated pathways were pyrrolo-quinoline quinone (PQQ) proteins, tetrapyrrole methyltransferases, and ABC transporters. Only two genes had down-regulation of less than log_2_-fold change −2, and these were an iron transporter and lysine 6-monooxygenase.

**Figure 7:**
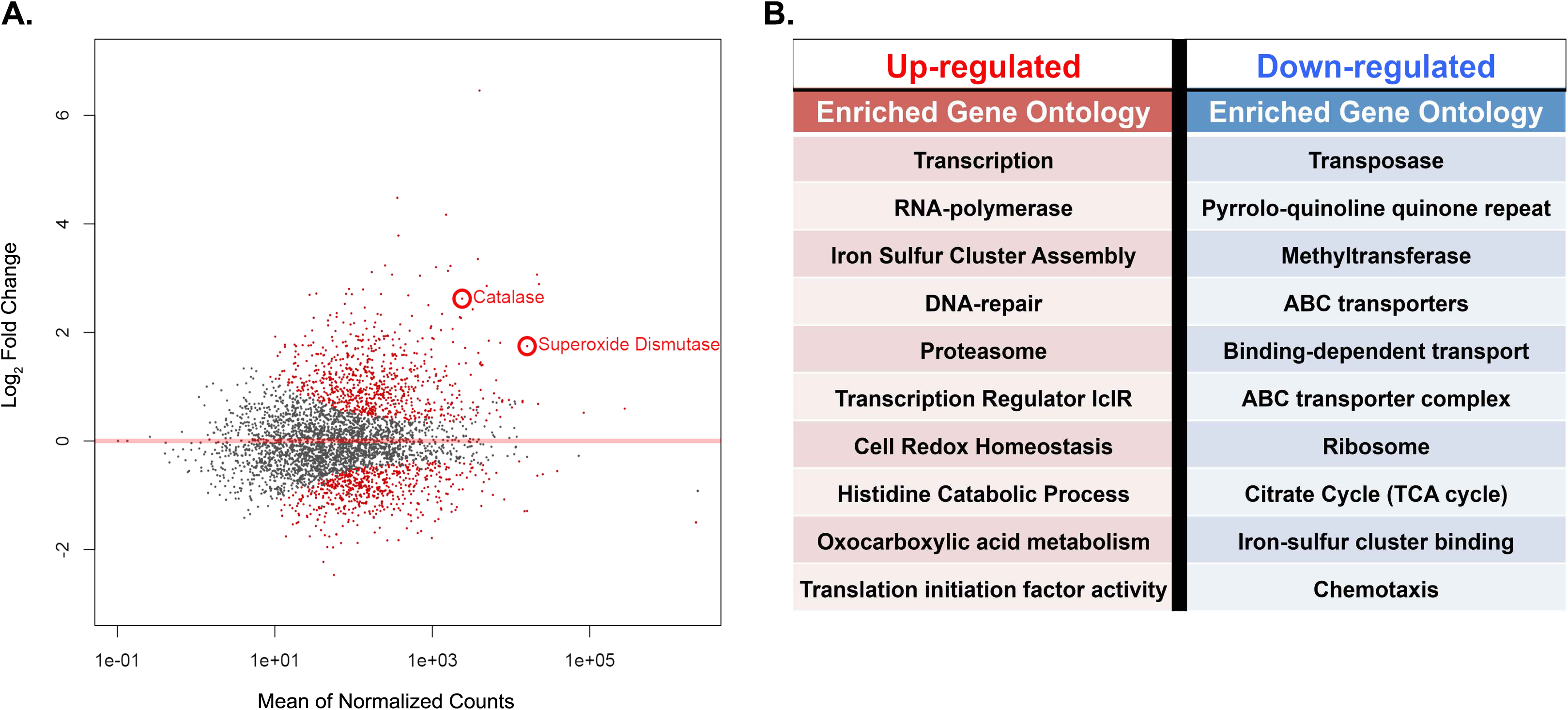
Distribution of differentially expressed genes during oxidative stress in *H. volcanii*. (A) MA-plot of differentially expressed genes; each point represent a gene. Significant (FDR < 5%) differentially expressed sRNAs are labeled in red. Circled points are known ROS scavengers. (B) Gene function for the most up- and down-regulated sRNAs.

## DISCUSSION

Previous studies of sRNAs in *Archaea* revealed the abundance of sRNAs within the third domain of life and have been pivotal in establishing a working hypothesis on archaeal sRNA functionality. These studies have been limited to (1) microarray studies that do not allow *de novo* discovery of sRNAs, (2) differential RNA-seq approaches (dRNA-seq), which selects only for primary transcripts and does not provide length (nt) information, nor expression information (only coverage), and (3) individual sRNAs studies, which do not give a holistic view of the pathways being regulated within the cell. Using a custom strand-specific sRNA-seq library preparation and analysis pipeline, we have developed a method to perform high-throughput analysis of sRNA transcriptional landscape, expression, regulatory effects, and to identify regulated gene pathways in response to environmental stressors within the *Archaea*. Through this study, we propose that sRNA-mediated transcriptional regulation is key in regulating stress responses to environmental challenges, such as oxidative stress, in the *Haloarchaea*. sRNAs have the potential to fine-tune the regulation of genes involved in the oxidative stress response resulting in increased resistance to extreme environmental stressors.

The discovery that sRNAs comprised nearly half the total transcriptome of *H. volcanii* and included basal transcriptional promoters, during both non-challenged and oxidative stress conditions, suggests that sRNAs have an important functional role under a variety of environmental conditions. We discovered thousands of sRNAs expressed in *H. volcanii* with the majority being antisense to genes, indicating that antisense transcription was ubiquitous within the cell. This is in stark contrast to most of the literature reporting that a majority of sRNAs discovered in *Archaea* were intergenic.(13, 22, 31, 36) This discrepancy is likely due to previous studies being limited to microarray approaches. Indeed, a recent study using directional RNA-seq (dRNA-seq) to map all transcription start sites (TSS) in *H. volcanii* found thousands of novel transcript TSS with 1,244 of these TSS being antisense to mRNAs.(24) Most of the TSS (75%) of the sRNAs we discovered in *H. volcanii* had the same TSS (+/− 5 nt) than those found in the dRNA-seq study by *Babski et al*, 2016 (24), further confirming our results. This underlines the importance of HTS studies, especially strand-specific RNA-seq such as our study, to discover the full extent of antisense sRNA expression in *Archaea*.

Our finding suggests that *cis*-acting sRNAs may play a larger role than *trans*-acting sRNAs within the cell but it should not be overlooked that the difficulty in finding *in silico* targets for intergenic sRNAs, because these sRNAs do not form 100% complementarity with their targets, might suggest that they have multiple mRNA targets. Antisense and intergenic sRNAs are broad classifications used in the archaeal small non-coding RNA field but our data revealed that further classification can be done based on sRNA-mRNA binding characteristics (5’ UTR, 3’ UTR, CDS), differential expression, and regulatory effects. We found that only a small fraction of antisense sRNAs targeted the 5’ UTR of mRNAs, which is in concurrence with work demonstrating that most mRNAs in *H. volcanii* are leaderless (lacking a 5’ UTR). However, further work is needed to determine whether translational repression via the masking of the Shine-Dalgarno sequence, as seen in bacterial sRNAs, is the mode of action of these 5’ UTR-binding sRNAs in *Archaea*.(37-39) A majority of the 5’ UTR-binding sRNAs targeted transposons, providing further evidence that they may constitute their own class of sRNAs. Within this context, 3’ UTR-binding sRNAs should also be considered another class of sRNAs, resembling eukaryotic sRNAs, and likely acting on the degradation of their target transcript. The majority of the antisense sRNAs we identified in *H. volcanii* had 100% complementarity within the CDS of their target mRNAs. This is the first report of such a finding in any domain of life and might constitute an attribute unique of archaeal sRNAs. We could not identified any ‘seed’ binding region for these CDS-binding sRNAs indicating that they likely have full occupancy upon the mRNA. It is also worth noting that there were only a few instances (<20 total) where CDS sRNAs overlapped more than the full length of the target mRNA or overlapped multiple *cis*-mRNA targets, which might also be unique to Archaea.

Most of *H. volcanii* sRNAs had a normalized expression value of 1 TPM or less meaning that sRNA transcripts were not abundant within the cell. In comparison, most mRNAs within *H. volcanii* had at least 20 TPM in expression value. Despite this order of magnitude difference between sRNA (low) and mRNA (high) expression levels, the top 5% of the most highly expressed sRNAs had higher expression compared to their mRNA target, suggesting a negative regulatory role in sRNA-mRNA interactions (**Fig 3b**).(40-42) This trend extended to a majority of sRNAs (both non-challenge and oxidative stress conditions) down to sRNAs with 1 TPM in expression level. Further evidence for a negative regulatory effect lies with up-regulated sRNAs. Most up-regulated sRNAs had target mRNAs that were down-regulated indicating sRNAs negatively regulate mRNA targets at the transcript level. Whether this negative regulation is occurring during transcription initiation/elongation or if up-regulated sRNAs are causing mRNA degradation is currently unknown. All the sRNAs targeting transposons at the 5’ UTR were up-regulated and the transposon mRNA down-regulated (**Fig 3b**), suggesting that these sRNAs might have a similar mechanistic function.(6, 43, 44) If indeed sRNAs are negatively regulating their target mRNAs in *H. volcanii*, we expected to find that down-regulated sRNAs have up-regulated target mRNAs. While some down-regulated sRNA targets exhibited this pattern, further supporting negative regulation, many mRNA targets were also down-regulated. Alternative hypotheses, reflecting the complexity of transcriptional regulation in the Archaea, can be formed: (1) some of these sRNAs may have a positive regulatory effect, such as stabilizing target mRNAs and masking them from degradation, (2) *trans*-acting intergenic sRNAs might be targeting these mRNAs, negatively regulating them, and (3) some may have an unknown function.(23) Despite this, more than twice the number of up-regulated antisense sRNAs (31) had a robust log-fold change (≥2) compared to down-regulated sRNAs (15) which suggests that up-regulating antisense sRNAs to down-regulate mRNA targets is the main strategy during oxidative stress.

The most enriched negatively regulated sRNA targets were transposases, chemotaxis proteins, and transcription factors. It has been demonstrated that transposons are opportunistic during stress conditions and can wreak havoc by hopping around in the genome causing double strand breaks, hence a need to be silenced.(6, 43, 44) A functional enrichment of IS4 transposon genes being down-regulated during oxidative stress supports our observation that up-regulated sRNAs negatively regulate transposons and suggests that transposon activity is tightly regulated during oxidative stress in *H. volcanii*. sRNA-mediated regulation of chemotaxis transducer proteins during oxidative stress suggests interesting implications in sensing ROS and motility. *H. volcanii* expresses a flagella homolog named ‘archaella’, which is organized into an operon and is regulated by a network of regulators called the archaellum regulatory network (arn) (identified in crenarchaea).(45, 46) The regulation of these motility genes is still under investigation and so far is restricted to a few examples such as H_2_/nitrogen limitation conditions in *M. janaschii and M. maripaludis*.(46-49) No direct transcriptional regulators of the archaellum have been identified in any euryarchaeota, but the deletion of archaellin genes, the presence of the H-domain set of type IV pillins, and agl proteins have been shown to affect the assembly of archaella in *H. volcanii*.(46, 50-52) Integral to how microorganisms maintain homeostasis in stressful and fluctuating environments are gene regulatory networks composed of interacting regulatory transcription factors and their target gene promoters.(53) Our discovery that sRNAs are targeting transcription factors provides evidence that sRNAs are likely deeply interlaced within complex gene regulatory networks of *H. volcanii* and these sRNAs are key to maintaining homeostasis during environmental stress such as oxidative stress. Many mRNA-targets of differentially regulated sRNAs were hypothetical proteins indicating that important genes in the oxidative stress response remain to be elucidated.

Our whole transcriptional analysis demonstrated that more than a quarter of the genes (~1100 – 30%) in *H. volcanii* were differentially regulated during constant H_2_O_2_-induced oxidative stress at ~80% survival, which is in agreement with the transcriptional response of *H. salinarum* during constant H_2_O_2_- (929 genes – 38%) and paraquat-induced (1099 genes – 45%) oxidative stress over 2 hours at ~80% survival.(2) This indicates that transcriptional regulation is crucial in order to mount this oxidative stress response via gene activation and repression. Two single-stranded DNA binding proteins (RpaB and RpaC) were found to be required for increased survival of *H. volcanii* to ionizing radiation (a proxy for desiccation) and UV radiation, stressors that both cause oxidative stress (54, 55) (DiRuggiero lab, data unpublished). In *H. salinarum*, Rpa operons were up-regulated during ionizing radiation as well and contributed to resistance.(56, 57) In conjunction to previous findings, we observed that two of the most up-regulated genes during H_2_O_2_ oxidative stress were RpaB and RpaC confirming their role in oxidative stress resistance in *H. volcanii* and likely other haloarchaea. One gene, a reactive intermediate/imine deaminase RidA-homolog, was up-regulated orders of magnitude more than any other gene. The encoded protein is known to be involved in synthesis of branched-chain amino acids by speeding up the IlvA-catalyzed deamination of threonine into 2-ketobutyrate.(58, 59) Previous work has shown that in the presence of reactive chlorine species (RCS), such as HOCl, imine deaminase seemed to inhibit IlvA activity suggesting that imine deaminase may have a different function in the presence of RCS.(58, 60) Further studies found that imine deaminase can sense RCS and in doing so becomes a chaperone that prevents protein aggregation.(58) Reactive oxygen species in hypersaline environments produce RCS.(61) In addition, ROS causes extensive, irreversible protein damage such as carbonylation, which in turn causes protein aggregation.(62, 63) This reactive intermediate/imine deaminase is the most up-regulated protein-encoding gene, suggesting that it may be playing a similar chaperon role to prevent protein aggregation, either sensing ROS or RCS produced by H_2_O_2_.(58, 60)

This is the first study to report on the transcriptional response of *H. volcanii* to oxidative stress and, while we found similar responses to H_2_O_2_ exposure than previously reported for *H. salinarum*.(2), further validating our work and providing evidence that *Haloarchaea* have evolved similar strategies to survive their environmental stresses, we also found responses that were unique to *H. volcanii.* Similarities to *H. salinarum* include the up-regulation of ROS scavenging proteins (catalase, superoxide dismutase), iron sulfur assembly proteins (SufB, SufD), proteasome genes, indicating high protein turn-over, and many DNA-repair genes.(2) Most of the down-regulated genes were involved with metabolism, such as sugar/phosphate/peptide ABC transporters, electron carriers (halocyanin), and TCA cycle enzymes, possibly to halt growth until damage is repaired.(2, 64). The most down regulated gene was an iron ABC transporter, most likely to limit further production of ROS via Fenton reactions.(2) Of unique responses to oxidative stress in *H. volcanii,* we found that all of the RNA polymerase subunits, transcription elongation factors, and transcription initiation factors were up-regulated in response to oxidative stress. The increase in sRNAs during oxidative stress could be attributed to this increase in transcription machinery. The majority of the 30S and 50S ribosomal subunits were down regulated, in contrast to *H. salinarum*. The up-regulation of histidine biosynthesis and catabolism into glutamate, and 2-Oxocarboxylic acid metabolism were unknown to be involved in the oxidative stress response, which further demonstrates there are still more mechanisms to uncover for oxidative stress resistance. RosR was identified as a global transcriptional regulator in *H. salinarum* and it strongly up-regulated during oxidative stress.(5) RosR demonstrated no differential expression to oxidative stress in *H. volcanii* indicating that it may be playing another role in this organism. Cell cycle genes (parA, cdc6) involved in chromosome segregation(65) were down regulated, further suggesting that division is being arrested (halting growth) in order to repair damage.

In this study, we showed for the first time that small non-coding RNAs are specifically associated with the oxidative stress response in Archaea. During oxidative stress, antisense sRNAs were prevalently transcribed, comprising nearly 30% of the transcriptome of *H. volcanii*, and most up-regulated antisense sRNAs imparted a negative regulatory effect on target mRNAs. These results support the hypothesis that antisense sRNAs in Archaea behave similarly to *cis*-acting bacterial sRNAs and eukaryotic siRNAs, which negatively regulate mRNAs by sharing extensive complementarity and facilitating RNA degradation.(66, 67). The precise mechanism(s) of sRNA-mRNA mediated regulation remains to be elucidated and in particular whether proteins are required to complex with sRNAs in order to mediate gene regulation such as in Bacteria (Hfq) and Eukarya (Ago). We also identified several classes of antisense sRNAs, based on their mRNA-binding patterns (3’ UTR, 5’ UTR, and CDS), and showed that CDS-targeting of mRNAs was the predominant mode of action for sRNA hybridization. Mechanistic differences between these classes of sRNA still need to be investigated as well as the regulatory roles of sRNAs in Archaea and their functional importance in adaption to extreme environments.

## MATERIAL AND METHODS

### Culture growth conditions

*H. volcanii* auxotrophic strain H53 (*Δ pyre2, Δ trpA*) was used for all experiments. Culturing in liquid and solid media was done in rich medium (Hv-YPC), at 42°C and with shaking at 220 rpm (Amerix Gyromax 737).(68) Uracil and tryptophan were added to a final concentration of 50 µg/mL, each.

### Oxidative stress exposure

We exposed 5 biological replicates of *H. volcanii* strain H53 liquid cultures to the oxidative stress agent H_2_O_2_. Initially, cultures were grown in 80 mL of Hv-YPC under optimal conditions to an OD of 0.4 (mid exponential phase). To ensure homogeneity, each replicate was subsequently split into two 40 mL cultures, one used for the non-challenged (control) condition and the other for the oxidative stress condition. For the latter condition, 2 mM H_2_O_2_ (80% survival rate, previously determined) was directly added to the cultures followed by an hour incubation at 42 °C with shaking at 220 rpm. Cultures were then rapidly cooled down, centrifuged at 5,000 x g for 5 minutes and the pellets resuspended in 18% sea water. The cell suspensions were then transferred to a 1 mL tube and centrifuged at 6,000 x g for 3 minutes, the pellets were flash frozen and stored at −80 °C until ready for RNA extraction. Control non-challenged culture replicates were processed in the same manner without the addition of H_2_O_2_.

### Oxidative stress survival curves

Assessment of survival in *H. volcanii* under oxidative stress conditions was done using microdilution plating as described before.(7) Counts were averaged and standard deviation calculated between replicates. Survival was calculated as the number of viable cells following H_2_O_2_ treatment divided by the number of viable untreated cells and graphed with standard error bars.

### RNA extraction

Total RNA was extracted using the Zymo Quick-RNA Miniprep kit with the following modifications: after addition of RNA lysis buffer to the frozen cell pellets, cells were processed with a 23 G needle and syringe to insure complete cell lysis. *H. volcanii* liquid culture is slimy and viscous thus to increase cellular lysis a 23 G needle and syringe was used to break down the cell pellet. Total RNA was then extracted following the standard kit protocol.

### Small RNA-sequencing library preparation (sRNA-seq)

Total RNA, for each biological replicate and condition, was size-selected using denaturing polyacrylamide gel electrophoresis. 20 µg of total RNA was loaded onto a 7% denaturing urea polyacrylamide gel (SequaGel, National Diagnostics) in 0.5 x TBE buffer and ran at constant power of 30 W until bromophenol blue bands reached the bottom of the gel. The gel was stained with SYBR Gold, visualized on a blue light box, and bands in the 50-500 nucleotide range, as indicated by the RNA Century Marker plus ladder (ThermoFisher), were excised. Small RNAs (sRNA) were eluted by rotating overnight in 1.2 mL 0.3 M NaCl, ethanol precipitated, and DNase I (NEB) treated (37 °C for 2 hours) as previously described.(69) Strand-specific libraries were prepared using the SMART-seq Ultralow RNA input kit (Takara), insert sizes checked with the Bioanalyzer RNA pico kit (Agilent), and paired-end sequencing (2 x 150 bp) was carried out on the Illumina HiSeq 2500 platform at the Johns Hopkins University Genetic Resources Core Facility (GRCF). Total RNA was rRNA-depleted using the Illumina Ribo-zero Bacteria kit. Library preparation and sequencing was as described above, omitting the size-selection by denaturing gel electrophoresis.

### sRNA- and RNA-seq read quality control and reference-based read mapping

Assessment of the quality of each sequencing library read was determined using fastqc. The program trim galore was used with base settings to trim adapter sequences from reads and to filter out low phred score reads (<20). Short length reads were preserved. Reads from each replicate were aggregated together per condition to get a set of consensus sRNAs and were mapped against the *H. volcanii* NCBI refseq genome (taxid 2246; 1 chromosome, 4 plasmids) using the hisat2 aligner with strand-specific options turned on and splice aware options turned off, paired-end mode.(70)

### sRNA- and RNA-seq transcriptome assembly

The reference-based alignments were assembled into transcriptomes using the program stringtie in order to build full-length transcripts, calculate coverage and expression values (TPM). The assembly was guided by a gene annotation file from the *H. volcanii DS2* (NCBI refseq taxid 2246) genome to build transcripts and annotate them either as a gene or novel transcript.(71) A minimum distance between reads for transcript assembly was specified at 30 nucleotides. gffcompare under default options was used to compare the assembled transcriptomes against the gene annotation file from *H. volcanii DS2* (NCBI refseq taxid 2246) to annotate transcripts as genes or non-coding RNA (antisense or intergenic).(72, 73) In house python scripts were used to bin transcripts that were annotated as genes, transcripts annotated as antisense (classified as non-coding region opposite from a coding region), transcripts annotated as intergenic (classified as non-coding region between two coding regions), and subsequently binned antisense sRNAs as 3’ UTR, 5’ UTR, or CDS overlapping.

### sRNA- and RNA-seq differential expression analysis

We used a read count-based differential expression analysis to identify differentially expressed sRNAs during oxidative stress. The program featureCounts was used to rapidly count reads that map to the assembled sRNA transcripts (described above).(74) featureCounts was run with strand-specific options on, paired-end mode on, multi-mapping off.(74) The read counts were then used in the R differential expression software package DESeq2.(75) Briefly, read counts were converted into a data matrix and normalized by sequencing depth and geometric mean. Differential expression was calculated by finding the difference in read counts between the non-challenged state (control) normalized read counts from the oxidative stress normalized read counts.(75) The differentially expressed sRNAs were filtered based on the statistical parameter of False Discovery Rate (FDR) and those that were equal to or under a FDR of 5% were classified as true differentially expressed sRNAs.

### in silico validation of sRNAs

Differentially expressed sRNAs were validated by two *in silico* methods: 1) Visualization of transcripts, and 2) open reading frame protein homology search. In the first method, transcriptomes for each replicate and condition were visualized on the Integrated Genome Viewer (IGV) against the *H, volcanii* (NCBI refseq taxid 2246) genome and annotation.(76) The sRNA transcript coordinates were used to locate putative sRNAs and if it was found within an operon it was eliminated from further analysis. In the second method, blastx (default parameters) was used to search for protein and domain homology for each sRNA and those that had significant homology with known proteins or domains were eliminated from further analysis.(77)

### Regulatory element motif identification of sRNAs

100 nucleotides upstream and downstream from the sRNA transcript start and stop coordinates were extracted using in house python scripts. These regions were searched for transcription motifs (BRE, TATA-box) using both multiple sequence alignments and visualization with WebLogo (default parameters) and motif searching with MEME and CentriMo (default parameters).(78, 79)

### In vivo validation of sRNAs by Northern Blot analysis

20 µg of total RNA and P^32^ ATP end-labeled Century+ RNA markers were loaded onto 5% denaturing urea polyacrylamide gels (SequaGel, National Diagnostics) and run at 30 watts for 1.5 hours to ensure well-spaced gel migration from 50 to 1,000 nucleotides (nt). Gels were transferred onto Ultra-hyb Nylon membranes and hybridized with 2 types of probes. For lowly expressed sRNAs, we probed with [γ-P^32^]dATP randomly primed amplicons generated with custom primers based on sRNA transcript genomic coordinates as determined by the sRNA-seq *in silico* analysis. Probe primers were at a minimum 10 nt inwards from the predicted genomic coordinates (start and stop) to ensure accurate transcript detection. Hybridizations were done at 65°C. To determine strandedness of sRNAs, we used [α-P^32^]dATP end-labeled oligo probes (20-23 nt) that were antisense to sRNAs. Hybridizations were at 42°C. The rpl30 protein (HVO_RS16975) transcript was used as a loading control for differential expression calculation because it was not differentially expressed under oxidative stress in this RNA-seq dataset. Differential expression was calculated using ImageJ.

### Gene Ontology (GO) enrichment analysis of mRNA-targets

NCBI gene names for all mRNA-targets of antisense sRNAs were uploaded into **D**atabase for **A**nnotation, **V**isualization and **I**ntegrated Discovery (DAVID) to determine the pathways and gene ontologies targeted by sRNAs.

### RNA-seq data

All raw read and processed data from these experiments are available at NCBI under BioProject PRJNA407425. Illumina raw sequence data (.fastq) for each replicate and condition are deposited in NCBI Sequence Read Archive with accession number SRP117726.

## ACKNOWLEDGEMENT

This work was supported by grant FA9950-14-1-0118 from the AFOSR.

We thank Madeline Cassani, Dr. Zhao Zhang, and German Uritskiy for advice and guidance on sRNA-seq library preparation, Evan Hass and Dr. Vidya Balagopal for advice on northern blotting, Dr. Jacques Ravel, Mike Humphrys, and David Mohr for sequencing efforts and technical advice, and Dr. John Kim and Dr. Sarah Woodson for helpful discussions.

